# Stressed Overwintering Bottleneck Hypothesis: Ocean warming and acidification synergistically disrupt Arctic zooplankton overwintering

**DOI:** 10.1101/2025.11.17.688696

**Authors:** Jildou Dijkstra, Luise Schott, Nele Thomsen, Helena Reinardy, Mathieu Lutier, Janne E. Søreide, Khuong V. Dinh

**Affiliations:** Freshwater and Marine Ecology (FAME), University of Amsterdam, P.O. Box 94248, 1090 GE Amsterdam, The Netherlands; The University Centre in Svalbard (UNIS), P.O. Box 156, Longyearbyen, Norway; Freie Universität Berlin, Institut für Biologie, Altensteinstr. 6, D-14195, Berlin, Germany; Scottish Association for Marine Science, PA37 1QA, Oban, Scotland; Section for Aquatic Biology and Toxicology, Department of Biosciences, University of Oslo, Blindernveien 31, 0371 Oslo, Norway

**Keywords:** Calanus, Climate Change, Diapause, Ocean warming, Ocean acidification, Polar Ecosystems, Svalbard

## Abstract

Ocean warming (OW), driven by the influx of warm Atlantic water masses, and acidification (OA) are threatening Arctic marine ecosystems. However, their potential synergistic effects are poorly understood, especially during the Polar Night when marine species are particularly vulnerable to stressors. Here, we tested our novel Stressed Overwintering Bottleneck Hypothesis (SOBH): warming will disrupt the overwintering of the keystone pan-Arctic copepod *Calanus glacialis,* a pivotal secondary producer, by impairing fitness-related traits underpinning survival and reproduction. We exposed *C. glacialis* to current and projected future OW levels (0 °C and 4 °C) and OA levels (pH 8.0 and 7.4-7.3) for 53 days during the mid-Arctic Polar Night. We assessed survival, development, and physiological and molecular mechanisms (oxygen consumption, lipid depletion, the expression of nine targeted genes related to oxidative stress and damage repair, and DNA damage). OW alone did not affect *C. glacialis* mortality; however, OA increased copepod survival at 0 °C. Notably, their combined effects (OWA) synergistically doubled mortality, as predicted by SOBH. Warming also accelerated moulting from copepodite stage V to adulthood in December, and increased respiration, exhausted lipid reserves entirely by early March, approximately one to four months before the spring algal bloom, further supporting SOBH. DNA damage and gene expression patterns indicated low investment in maintenance and damage repair. Collectively, these findings reveal hidden mechanisms by which OW and OA synergistically threaten overwintering *Calanus* copepods by drastically increasing mortality, accelerating moulting, raising metabolic rates, and causing early lipid depletion. These effects generate cross-seasonal phenological mismatches among overwintering survival, energy reserves, reproduction, and primary production. Such stressed overwintering bottlenecks in foundational secondary producers like *Calanus* copepods provide novel explanations for how OW and OA can constrict Arctic marine food webs. At a broader perspective, SOBH highlights how multiple stressors induced overwintering disruption could reshape pan-Arctic and global biodiversity.

## 1. INTRODUCTION

Arctic marine ecosystems face unprecedented challenges due to rapid environmental changes driven by anthropogenic climate change (IPCC, 2021). Atmospheric CO_2_ levels are increasing at a rate unmatched in the last 300 million years (Hönisch et al., 2012), resulting in ocean warming (OW, increasing seawater temperatures) and acidification (OA, decreasing seawater pH, IPCC, 2021). The OW and OA rates in Arctic seas are three to four times faster than in other seas and oceans (Qi et al., 2022; Rantanen et al., 2022; Årthun et al., 2025). OA has crossed planetary boundaries (Findlay et al., 2025). These accelerated changes pose severe threats to Arctic marine species, which are typically adapted to a narrow range of low temperatures (e.g., Carstensen et al., 2012; Dahlke et al., 2018). For example, OA is projected to reduce suitable habitats for polar invertebrates by up 63% (Findlay et al., 2025). A global analysis further indicates that within 50 years, marine zooplankton face compounded risks from both OW and OA which are projected to double compared to current levels, although data from Polar Night remain notably scarce (Richon et al., 2024).

The Polar Night, a period of prolonged darkness and low temperatures, is a critical yet understudied phase for overwintering Arctic marine organisms, during which energy conservation through lowered metabolism is essential for survival, moulting, gonad maturation, mating, and successful reproduction post-winter (Berge et al., 2020; Toxværd et al., 2018). Overwintering is, therefore, expected to increase the vulnerability to stressors (Berge et al., 2020; Toxværd et al., 2018). For example, a short-term (7-day) winter exposure of the Arctic pteropod *Limacina helicina* to warming increased the negative effect of ocean acidification on the shell degradation (Lischka & Riebesell, 2012). Winter OW, driven by the influx of warm Atlantic water masses in the Arctic marine ecosystems (Skagseth et al., 2020; Årthun et al., 2025) can disrupt overwintering through increasing metabolic rates of overwintering zooplankton that could result in depleting lipid reserves, and impairing gonad maturation (Karlsson & Søreide, 2024). Such overwintering disruptions can reduce survival, alter reproductive success, and cause phenological mismatches with primary production, potentially triggering cascading effects across Arctic ecosystems and global carbon cycles (Berge et al., 2020; Dinh et al., 2023; Jonasdottir et al., 2015; Oziel et al., 2025; Pinti et al., 2023; Søreide et al., 2010). These findings underscore the urgent need to investigate the combined impacts of OW and OA during this prolonged critical period in Arctic marine ecosystems, yet the interactive effects of these stressors on overwintering Arctic marine species remain largely understudied (but see Lischka & Riebesell, 2012).

In the Arctic marine ecosystems, the relatively large calanoid copepods of the genus *Calanus* comprise up to 70-80% of the zooplankton biomass (Daase & Eiane, 2007; Kosobokova & Hopcroft, 2010). These species are pivotal grazers on phytoplankton blooms, the primary energy source in lipid-based Arctic marine ecosystem (Beaugrand et al., 2003; Bouchard & Fortier, 2020; Leu et al., 2015; Søreide et al., 2010) and play a major role in transporting carbon to the deep ocean (Jonasdottir et al., 2015; Pinti et al., 2023). *Calanus glacialis*, a dominant Arctic species, has a wide pan-Arctic distribution, particularly abundant in the shelf seas and increasingly prevalent in the central Arctic Ocean (Ershova et al., 2021; Kvile et al., 2018). These foundational grazers have evolved a specialised life history to survive the long Polar Night and capitalise on the brief pulse of spring algal blooms, which are crucial for development, lipid accumulation, gonad maturation, and reproduction (Baumgartner & Tarrant, 2017; Leu et al., 2015; Søreide et al., 2010). Following the peak productivity phase, *C. glacialis* migrates to deeper, colder waters in winter, where it overwinters and is protected from predators (Daase & Søreide, 2021).

The present study addresses these knowledge gaps by examining how current and projected future OW and OA interact to disrupt the overwintering of *Calanus glacialis,* a key Arctic grazer, during the Polar Night. We propose and test **a novel hypothesis, “Stressed Overwintering Bottleneck Hypothesis (SOBH)”,** that winter warming will disrupt the overwintering of *C. glacialis* by increasing metabolism, depleting the lipid energy reserve, accelerating the development by moulting from copepodite V to adults, shortening the diapause period, and reducing survival. Overall, it will result in a mismatch in their phenology with the spring algal bloom and reproduction; all these important biological processes are enhanced by ocean acidification (Dahlke et al., 2018; 2017; Dam et al., 2021; Lischka & Riebesell, 2012). This SOBH can be applied to any stressor beyond OW and OA (e.g., chemical pollutants, light pollution, freshening, hypoxia) that can disrupt the overwintering of organisms across ecosystems spanning from the polar to subtropical regions (Dinh et al., 2023). To obtain mechanistic insights underlying the disruption of copepod overwintering, we analysed the DNA damage and the expression of genes relating to oxidative stress responses and the DNA repair system (Manandhar et al., 2015; Schieber & Chandel, 2014). Understanding these mechanisms in *C. glacialis* is critical to predict how multiple stressors interact to affect physiological processes, fitness, and population dynamics (Dinh et al., 2022; Schäfer & Piggott, 2018). To increase the realism of predicting the interactive effects of warming and OA, we exposed *C. glacialis* to these two stressors over 53 days in the middle of the Arctic Polar Night between December and January with multiple sampling times, thereby providing critical insights into biological processes and enabling us to model multi-level biological effects until the reproductive period and estimate the phenological mismatch with the Arctic algal blooms.

## 2. METHODS

### 2.1 Sampling

Sampling was conducted in Billefjorden (78°40’N;16°40’E, Figure 1), a seasonally ice-covered sill fjord on the western coast of Svalbard. Due to its two shallow sills (80m and 50 m) and location inside the North-eastern extremity of the Isfjord system, external water masses from the West Spitsbergen Current are prevented from entering, and cold (lower than -1°C) winter-cooled water persists in this glacial basin year-round below 80 m depth, supporting an abundant Arctic species-dominated zooplankton community (Søreide et al., 2022). Zooplankton were collected on the 12th of November 2022 by triplicate vertical WP-2 plankton net hauls (Hydro-Bios, area = 0.25 m^2^, mesh size = 180 µm). The net was hauled from 10 meters above the bottom depth (-185 m) and closed at 100 meters depth to target the overwintering *C. glacialis* population. Samples were stored in a black container at in situ sea temperatures. Within 8 hours of capture, the sample was diluted with surface seawater and kept at 0°C in the laboratory. *Calanus glacialis* copepodites at stage V (CV) were picked by eye based on estimated PL (> 3 mm) and red antennae colouring (Choquet et al., 2018; Nielsen et al., 2014). All sorted individuals were kept in seawater filtered through a 0.2 µm filter (FSW, Whatman, GF/F, diameter 47 mm, 2 µm pore) at 0°C before the experiment. No food was provided before or during the experiment, resembling natural food-restricted overwintering conditions. To confirm visual selection accuracy, 100 *Calanus* spp. individuals were subsampled at random from the net sample. These were visually identified as either *C. glacialis* or *C. finmarchicus* and stored at -80°C for molecular species identification following the methods described by Smolina et al. (2015). Those visually identified as *C. glacialis* and those visually identified as *C. finmarchicus* were confirmed to be 100% correctly identified with molecular identification.

**Figure 1.**
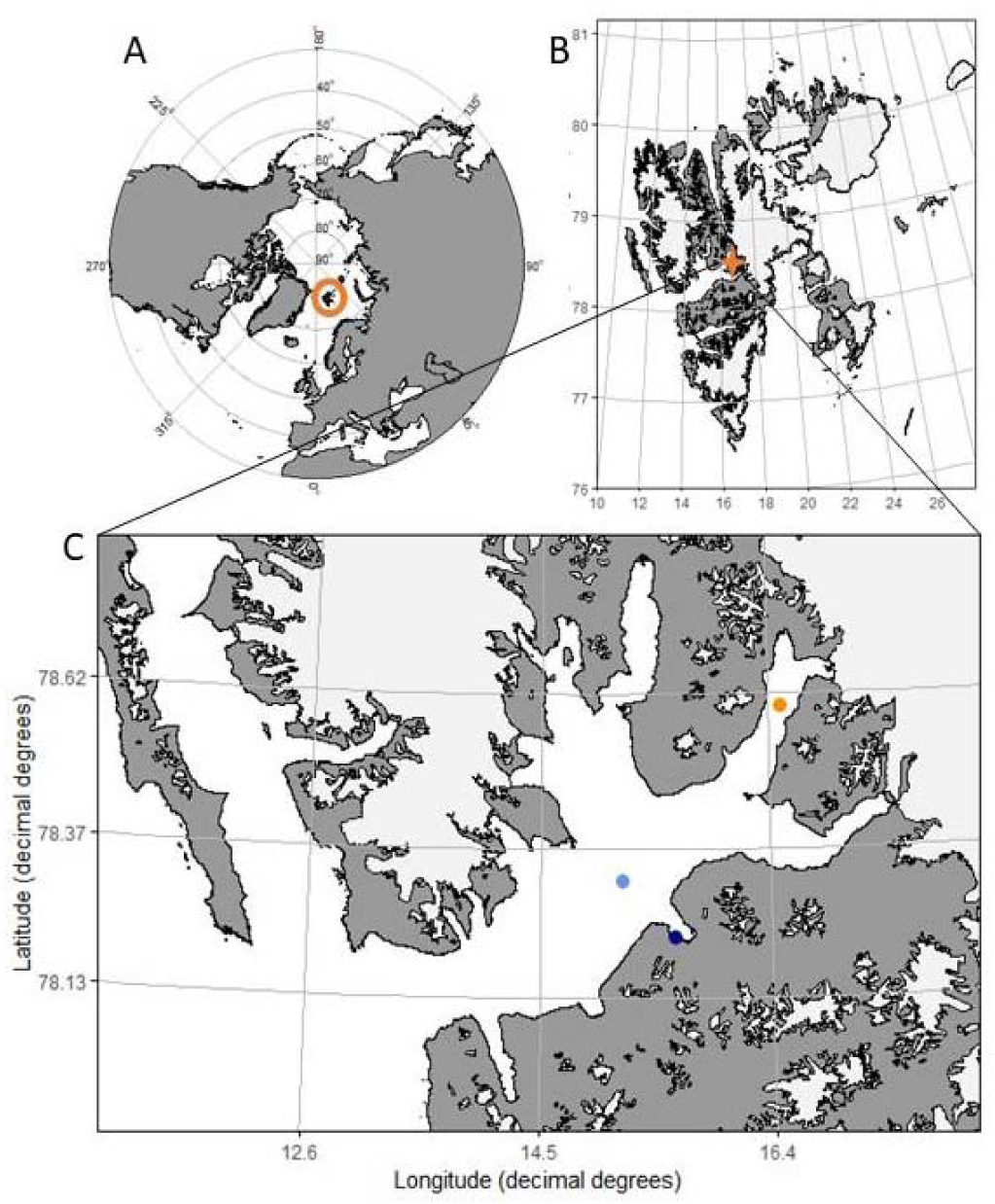
A map of (A) The location of the Svalbard archipelago is indicated by the orange circle (B) The location of the Isfjorden system on the Western coast of Svalbard, with Billefjorden is indicated by the orange star (C) The location of the Billefjorden sampling station (BAB) is indicated with the orange point (78°40’ N, 16°40’ E) and other water sampling locations indicated by the light blue point (ISK: 78° 18’91.2227” N and 015° 10’64.4063” E) and dark blue point (Harbour: 78° 13’ 45.1668” N, and 15° 36’ 3.9558” E).

### 2.2 Sea water treatments

The copepods were incubated in natural well-mixed sea surface waters due to cooling and ongoing winter convection, collected from the Billefjorden sampling station (BAB), the Isfjorden sampling station (ISK), and the Longyearbyen Harbour (Figure 1). All water was filtered through 0.7 µm and 0.2 µm GF/F glass microfiber filters (Whatman (R), Florham Park, NJ, USA) consecutively. After which it was stored at 4 °C. Seawater was oxygenated to 100% saturation using air bubbling before the experiment. Water was placed into the respective temperature treatment incubators 12 hours before use to adjust the water temperature. The pH of the lowered pH treatment water was adjusted just before use by adding CO2-saturated water (∼ 40 ml/5L) until the desired pH was reached. The pH for the nominal pH 8.0 treatment was left unadjusted and was the ambient pH of the natural seawater (about 8.0). The temperature, conductivity, and pH of all treatment water were measured and recorded before use. Temperature was measured using a temperature probe (Ebro TFX 410) with a fixed probe (Pt 1000), and conductivity was determined with a conductivity electrode (TetraCon 325-3) connected to a conductivity meter (WTW Cond 3210). The pH was measured using a pH probe (Sentix 41) connected to a pH meter (WTW pH 3110). The condition of the pH probe was checked before every water change using Certipur ® Merck NBS buffers (pH = 4.00 and 7.00) and calibrated on a total pH scale using a tris(hydroxymethyl)aminomethane (TRIS) buffer solution in Synthetic Seawater (salinity 35; obtained through A. Dickson CO2 QC Lab, Scripps Institution of Oceanography, University of California, San Diego, USA) once a week. Water samples (100 ml) from all treatments were obtained four times throughout the experiment (Days 20, 25, 40, and 45) for total alkalinity (A_T_) analyses. The sample was filtered through 0.7 µm (Whatman, GF/F, diameter 47 mm, 7 µm pore) and immediately fixed with 50 µl of 0.05% saturated mercury chloride solution and stored in the dark until analysis.

Dissolved inorganic carbon (DIC) measurements were performed at NIVA (Oslo, Norway), which is accredited for this type of analysis. The measured total pH (pH_T_), temperature, salinity, and DIC were then used to calculate total alkalinity (A_T_), partial pressure of CO_2_ (pCO_2_), and the saturation states of calcite (Ω_C_) and aragonite (Ω_A_) using the *seacarb* package in R (v4.2.1). Detailed results are provided in Table 1.

**Table 1.** Seawater carbonate chemistry treatment parameters. Total pH (pH_T_), T (°C), and S (PSU) were measured during each water change and averaged for the entire experimental period. The dissolved inorganic carbon concentration (DIC) was measured on days 20, 25, 40, and 45 throughout the experiment (N per treatment = 10). Partial pressure of CO_2_ (pCO_2_ in μatm), total alkalinity (A_T_), and the saturation states of calcite (Ω_C_) and aragonite (Ω_A_) were calculated using pHT, DIC, T, and S using the *seacarb* package in R. Values are depicted as the means ± s.d.

| Treatment | Nominal pH | Measured |  |  |  | Calculated |  |  |  |
| --- | --- | --- | --- | --- | --- | --- | --- | --- | --- |
|  |  | pH<sub>T</sub> | DIC<br>(µmol kg<sup>-1</sup>) | T (°C) | S (PSU) | P CO<sub>2</sub><br>(µatm) | A<sub>T</sub><br>(µmol kg<sup>-1</sup>) | Ω<sub>C</sub> | Ω<sub>A</sub> |
| Control | 8 | 7.9±0.1 | 2209±80 | 0.5±0.5 | 32.4±0.3 | 567±19 | 2269±83 | 1.57±0.09 | 0.97±0.06 |
| OW | 8 | 7.8±0.1 | 2264±21 | 4.1±0.5 | 32.3±0.2 | 742±10 | 2315±24 | 1.49±0.05 | 0.94±0.03 |
| OA | 7.5 | 7.4±0.1 | 2367±67 | 0.3±0.3 | 32.3±0.3 | 1881±53 | 2279±66 | 0.52±0.03 | 0.32±0.02 |
| OWA | 7.5 | 7.3±0.1 | 2388±22 | 4.1±0.3 | 32.2±0.2 | 2430±22 | 2286±21 | 0.49±0.01 | 0.31±0.01 |

### 2.3 Multiple stressor experiment

To test how OW may interact with OA to disrupt the overwintering of *C. glacialis*, 200 copepodite CV stage were incubated for each of the four treatments: 2 temperatures (0 and 4 °C) × 2 pH levels (8.0 and 7.5), resulting in 4 experimental treatments: control = 0 °C and pH 8, ocean warming (OW) = 4 °C and pH 8; ocean acidification = 0 °C and pH 7.5 and ocean warming and acidification (OWA) = 4 °C and pH 7.5. Actual pH levels were 7.9 ± 0.1 and 7.4 ± 0.1 at 0 °C and 7.8 ± 0.1 and 7.3 ± 0.1 at 4 °C (see Table 1). A +4°C warming and -0.5 to -0.6 pH unit fall within the projected future temperature and pH conditions in the Arctic seas and oceans by 2100 (IPCC, 2021; Årthun et al., 2025), especially in the Arctic fjord systems (Koziorowska-Makuch et al., 2023). These levels have often been used to assess the impacts of climate change on marine species (Brennan et al., 2022; Dam et al., 2021), including the Arctic zooplankton (Hildebrandt et al., 2016). The copepods were distributed amongst 20 bottles for each treatment (n = 10 individuals per bottle, n = 20 bottles per treatment) (Biological oxygen demand bottles 250 mL, VWR, Canlab or Duran Schott 250 mL, Duran, USA). Three bottles per treatment without copepods served as controls for any bacterial consumption or other oxygen fluctuations. To start the experiment, copepods in the warming treatment were warmed at a rate of 1°C per 4 hours just before the onset of the experiment inside the incubator (Termaks Cabinets-KB series, Nordic Labtech, Kungsbacka, Sweden). The temperature and light exposure inside each incubator were logged every 5 minutes using a HOBO® Pendant (Onset, Bourne, USA) temperature and light logger (UA-002-08) inside each incubator (Figure S1 and S2 in Appendix S1).

The copepods were incubated for a maximum of 53 days (December 1st, 2022 – January 22nd, 2023) as depicted by the experimental plan in Figure 2a. On days 10, 20, and 40 of the experiment, two bottles from each treatment were terminated for further analyses. Halfway through the experiment on day 28, 6 bottles per treatment were terminated, and 10 bottles from each treatment were terminated at the end of the experiment on day 53. At each of these sampling moments, individuals from half of the bottles were used to determine the lipid content, developmental stage, and dry weight, and the other half of the bottles were used for molecular analyses to determine DNA damage and gene expression.

**Figure 2.**
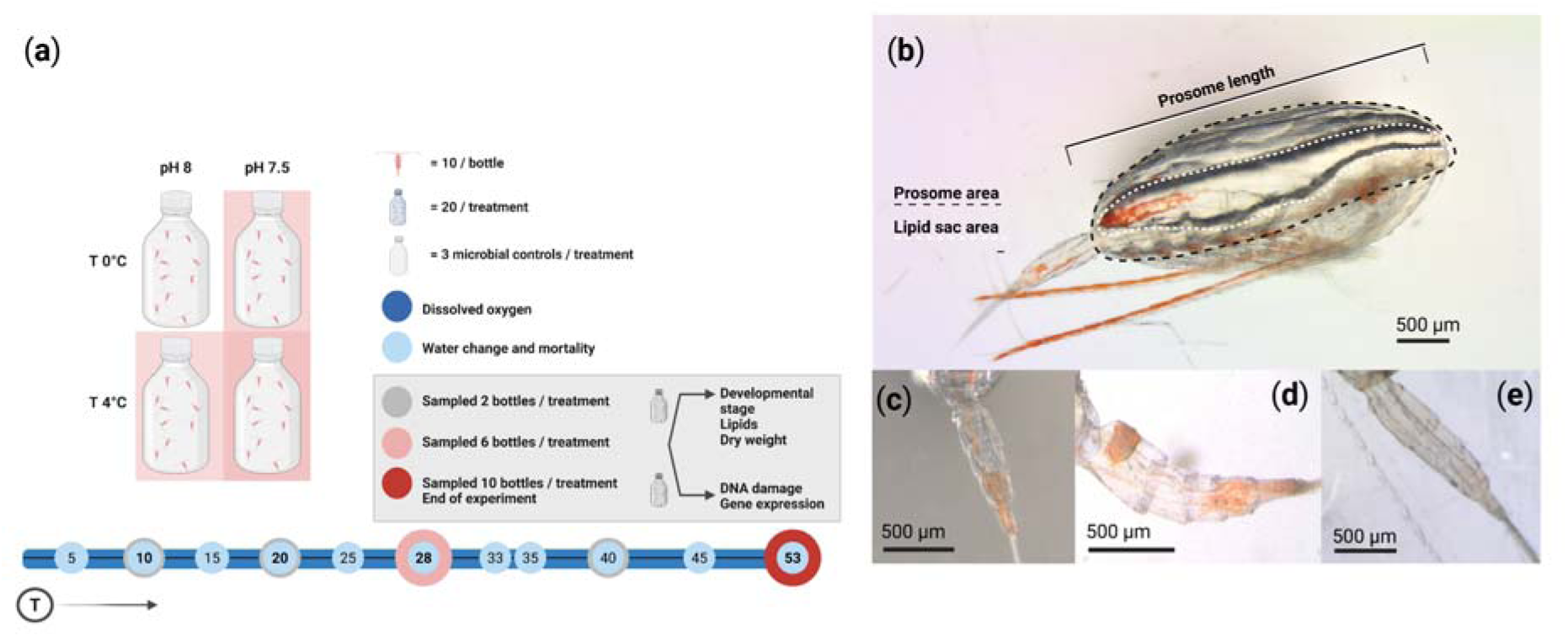
(**a**) Overview of experimental plan displaying treatments and replicates (top) and a timeline of sampling moments and purposes throughout the entire incubation period (bottom). (**b**) Ventral view of *Calanus glacialis* with prosome length, area, and lipid-sac area indicated. (**c**) Fifth copepodite stage urosome with 4 distinct segments visible. (**d**) Adult female urosome with 4 segments, of which the first one is visibly enlarged. (**e**) Adult male urosome with 5 distinct segments

To keep oxygen levels sufficiently high and maintain stable pH conditions, the incubation water was replaced every five days. Between days 45 and 53, the water was changed only once due to sustained high oxygen concentrations as observed by the daily dissolved oxygen concentration measurements. During each water exchange, the conductivity, temperature, and pH were measured inside each bottle to monitor steady exposure values (Table 1). Additionally, all individuals were inspected, and dead individuals were identified by developmental stage, measured for size, and subsequently removed.

### 2.4 Lipids, developmental stage, and dry weight

To determine the lipid content and developmental stage, individual images were taken from the ventral side using a Leica M205 C (Leica Application Suite version 4.12) or Leica MZ16 stereomicroscope fitted with a Leica MC170HD camera. The prosome length (PL), prosome area (PA), lipid-sac area (LA), and developmental stages were determined as depicted in Figure 2b-e, using ImageJ software (v1.54d). All individuals included in the experiment were stage CV at the onset of the incubation, so the presence of adult female or male urosome traits (Figure 2, c, d, e) was a direct result of individuals moulting into the final adult developmental stage.

The LA to PA ratio (LAPA) of each individual was calculated as a body size-dependent index for individual lipid content (Vogedes et al., 2010). The LAPA ratio is thus a measure of lipid fullness (Daase et al., 2018). To determine the lipid fullness at day 0, the LAPA ratios for 100 CV individuals from the original sampled pool of zooplankton were determined. After imaging, individuals were quickly dipped and rinsed in distilled water and placed in pre-weighed tin cups and then freeze-dried for 24 hours before being weighed again on a Microbalance XPR scale (accuracy ± 0.5 µg).

### 2.5 Oxygen consumption, gene expression, and DNA damage analyses

Twice daily dissolved oxygen levels were measured at 09:00 and 18:00, ensuring levels never dropped below 60% saturation throughout the entire incubation. Oxygen levels were determined using a fibre optic cable connected to the OXY-4 4-Channel Fiber-Optic Oxygen Meter (PreSensTM, GmbH, Regensburg, Germany) (Appendix S1).

Surviving zooplankton for DNA damage and gene expression analyses were fixed in 2 mL cryotubes in RNA*later* stabilization solution (Thermo Fisher Scientific, Waltham, MA, USA) (5:1 RNA*later*:FSW) immediately after being removed from the experimental bottle. Individuals were stored at -80°C for one month before transportation on dry ice to the Scottish Association for Marine Science (SAMS) for further analysis. RNA was extracted with the Qiagen RNeasy Mini Kit following the manufacturers protocol with adjustments for chitinous copepods (see Appendix S1).

The fast micromethod assay (FMM) for the detection of DNA strand breaks was conducted on individual copepods to quantify DNA damage. The method described by Halsband et al. (2021) was followed with adjustments for detection in individual copepods (see Appendix S1).

Nine genes were selected for their known function (Table S1) in either the oxidative stress response (*ferritin* and *cat*) or DNA repair system (*ercc1*, *ercc4*, *pcna*, *parp1*, *apex*, *ogg1*, *rad51*). Primers were extracted from literature or designed from published transcriptome sequences (Table S1). Primers were designed for a product length of 100 – 150 bp and checked for hairpin structures, complementarity, secondary structures, and self-annealing with Primer3Plus (Untergasser et al., 2012) and BLAST from the National Center for Biotechnology Information (NCBI) (see Appendix S1 for the detailed method).

### 2.6 Statistical analyses

All statistical analyses were done using R Statistical Software (v4.2.1) with RStudio (v2023.03.1+446). Packages used were ggplot2, ggpubr, dplyr, car, tidyverse, and ggoceanmaps. Statistical significance of the temperature and pH on mortality, LAPA ratio, moulting, and oxygen consumption was tested using multiple linear regression models (MLRM) in which both individual and combined effects of pH and temperature were analysed. Linear regression results were considered significant if the slope was significantly different from zero according to the t-statistic with a *P*-value < 0.05. Lipid trajectories were predicted by fitting treatment-specific linear models, allowing a small backward extrapolation to approximate realistic baseline reserves under continuous exposure, followed by forward projections until depletion. The interactive effect of temperature and pH on the oxygen consumption was tested using two-way analysis of variance (ANOVA) paired with a *post hoc* Tukey Honest Significant Difference test. To assess within time point differences in DNA damage (Strand Scission Factor, SSF) and gene expression, a one-way analysis of variance (ANOVA) with post hoc multiple range test was used to assess significant differences between treatment means with Fisher’s least significant difference (LSD) procedure at the 95% confidence level. Results were considered significant if the *P*-value < 0.05. All results are presented as a mean ± standard errors.

When the interactive effects of OW and OA are significant, we calculated the interaction effect size (estimated as Hedges’ d) with 95% confidence interval (CI) following the approach by (Dinh et al., 2016; Jackson et al., 2016). Hedges’ d values exceeding zero denote amplified, synergistic effects of OW and OA.

## 3. RESULTS

### 3.1 Mortality

Overall, mortality was low and lowest in the control (pH=7.9, 0 °C). Mortality increased over time; however, temperature alone had no effect on survival (Table 2, Figure 3a). OA interacted with temperature in two contrasting ways: (1) OA reduced the mortality rate at 0 °C, but (2) increased the mortality rate at 4 °C (OWA), generating a main positive effect of OA, and an OWA synergistic interaction (Hedges’ d = 0.24, 95% CI: [0.05 – 0.43], Table 2, Figure 3a).

**Figure 3.**
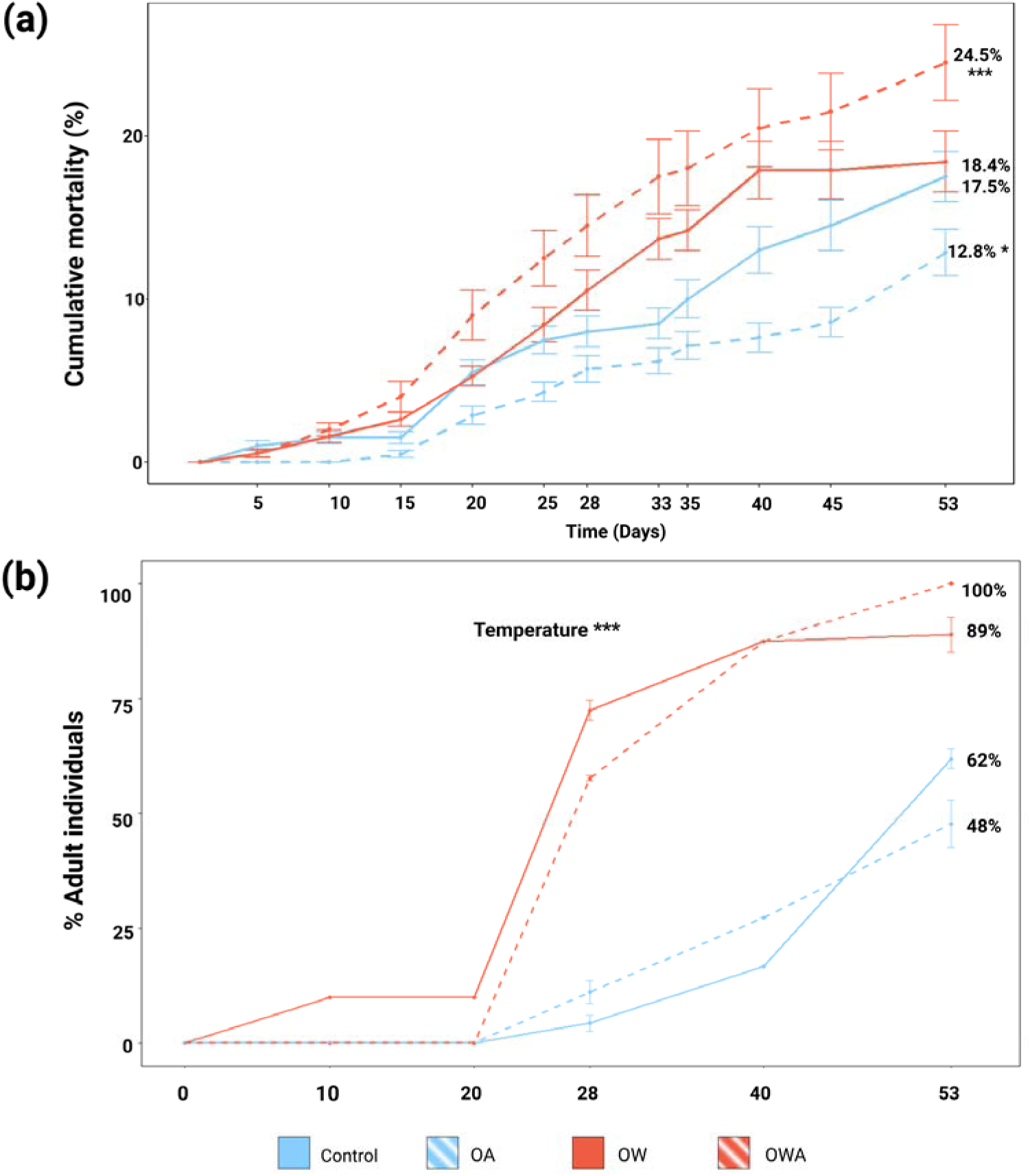
Mortality (**a**) and development (**b**) of *Calanus glacialis* in response to winter ocean warming (OW), acidification (OA), and their combined effects (OWA). The points represent the mean cumulative mortality bottle^-1^ treatment^-1^ ± standard errors (**a**), or the mean percentage of adult individuals bottle^-1^ treatment^-1^ ± standard errors (**b**). The red lines represent treatments exposed to a temperature of 4 °C, and the blue lines represent the colder treatments at 0 °C. The dotted lines indicate those treatments in which pH was lowered to 7.4-7.3. Asterisks * indicates significant influence on the slope throughout the entire 53 days of the experiment, MLRM, \**P* < 0.05 and \*\*\**P* < 0.001).

**Table 2.** Summary of the results of the Multiple Linear Regression (MLR) analyses for the effects of ocean warming (OW), acidification (OA) and their interactions (OWA) on mortality, and the Analyses of (co)variance (ANCOVA) for the effects of OW, OA and their interactions (OWA) on development, lipid content and oxygen consumption of *Calanus glacialis* following a 53-day exposure to single and combined effects of ocean warming and acidification.

| Mortality |  |  |  |  |  |  |
| --- | --- | --- | --- | --- | --- | --- |
| Factor | Model df | Estimate | Std. Error | t-value | *P* | Evidence |
| OW | 3 | 0.188 | 0.111 | 1.689 | 0.915 | No |
| **OA** | **3** | **-0.271** | **0.109** | **-2.496** | **0.013** | **Strong** |
| **OW × OA** | **3** | **0.549** | **0.155** | **3.536** | **0.0004** | **Very strong** |
| Development (Fraction moulted) |  |  |  |  |  |  |
| Factor | df | Sum Sq | F value | *P* | Evidence |  |
| **OW** | **1** | **1.6316** | **56.140** | **< 0.0001** | **Very strong** |  |
| OA | 1 | 0.0180 | 0.618 | 0.441 | No |  |
| OW × OA | 1 | 0.0096 | 0.329 | 0.572 | No |  |
| Residuals | 21 | 0.6103 |  |  |  |  |
| Lipid fitness (LAPA) |  |  |  |  |  |  |
| Factor | df | Sum Sq | F value | *P* | Evidence |  |
| **OW** | **1** | **0.1868** | **33.558** | **<0.0001** | **Very strong** |  |
| OA | 1 | 0.0052 | 0.938 | 0.346 | No |  |
| OW × OA | 1 | 0.000 | 2.122 | 0.123 | No |  |
| Residuals | 17 | 0.0946 |  |  |  |  |
| Oxygen consumption |  |  |  |  |  |  |
| Factor | df | Sum Sq | F value | *P* | Evidence |  |
| **OW** | **1** | **0.108** | **1.805** | **0.0014** | **Strong** |  |
| OA | 1 | 0.0059 | 0.9879 | 0.164 | No |  |
| **OW × OA** | **1** | **0.570** | **0.949** | **0.0019** | **Strong** |  |
| Residuals | 830 |  |  |  |  |  |

### 3.2 Development

Overall, OW accelerated copepodite development, with individuals moulting into adults approximately one month earlier than at 0 °C (main effect of Temperature, MLRM, Table 2, Figure 3b). Strikingly, OW triggered rapid moulting in two-thirds to three-quarters of the population in one week, between days 20 and 28 of the exposure period (20 – 28 December, mid-winter). In contrast, at 0 °C, no copepodites moulted within the first 20 days, and subsequent moulting occurred gradually without an abrupt change in the moulting rate (Figure 3b). By the end of the 53-day exposure period, approximately 95% of the individuals at 4 °C had moulted into adults, compared to only 55% at 0 °C. OA effect on moulting and its interaction with OW were not detected (Table 2).

### 3.3 Lipid fitness

The lipid fitness (or lipid/body condition) as indicated by the LAPA ratio decreased faster in both the warmer treatments (MLRM, *P*< 0.05), regardless of pH levels (Table 2, Figure 4a). Assuming that the LAPA reduction would be at similar rates for 0 and 4*°*C throughout the winter period, the lipid depletion (LAPA = 0) would occur on days 06, or 04 March (Table 3, Figure 4b), at least a month or longer before fresh algal food in the Arctic shelf seas around Svalbard and Greenland normally occur (Table S2, Appendix S2). In contrast, at 0°C, females could have lipid reserves available until the end of April (at OA) or June (control treatment), which would match the timing of the spring blooms (Table S2, Appendix S2).

**Figure 4.**
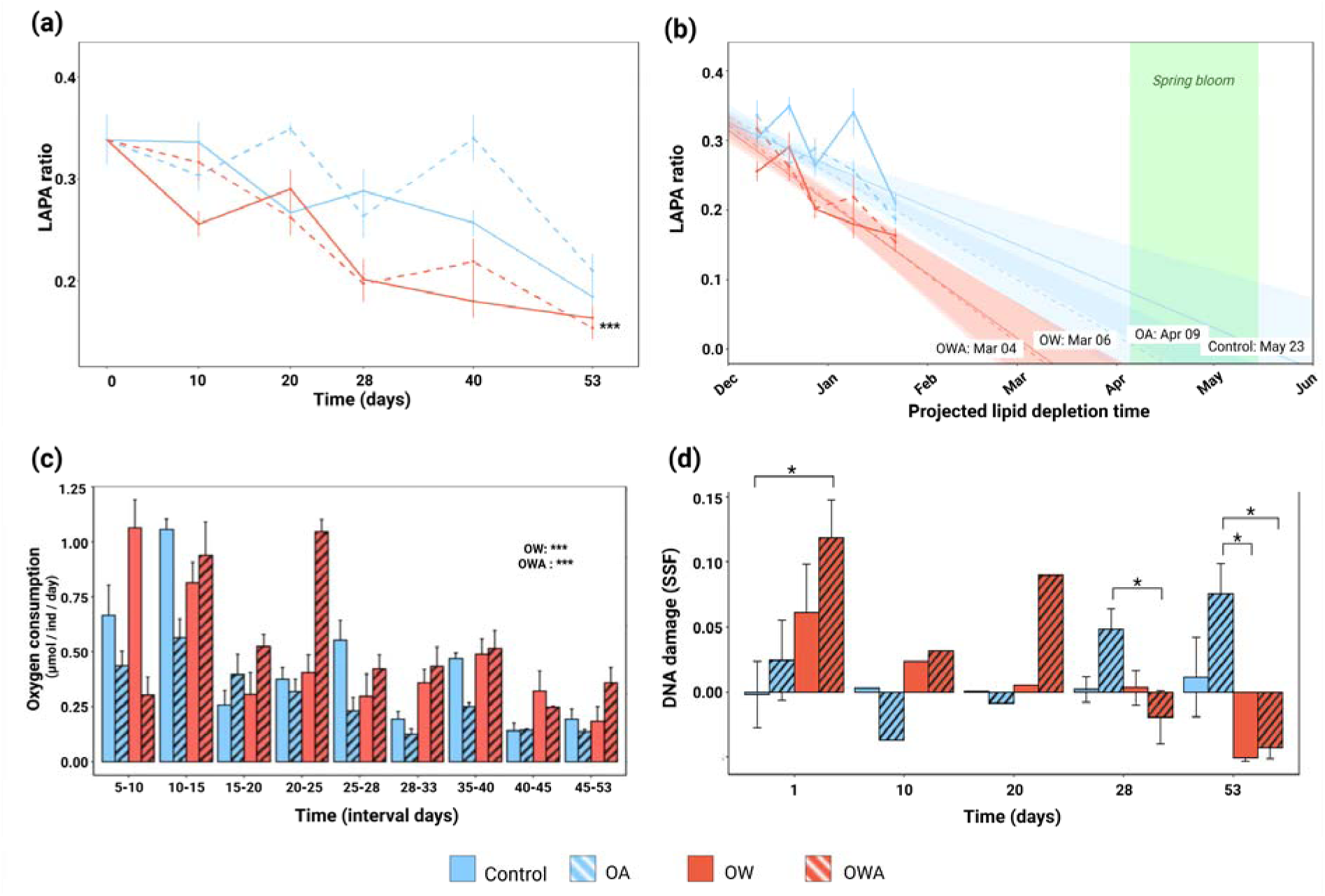
Lipid area to prosome area ratio of *Calanus glacialis* is depicted over the incubation period (**a**), and predicted time that the lipid would be completely depleted from linear regressions of each experimental treatment (**b**); (**c**) oxygen consumption rates in µmol individual^-1^ day^-1^ presented per week. The points represent the mean LAPA ratio for each treatment ± standard errors. (**d**) DNA damage (strand scission factor, SSF, fast micromethod) after acute exposure (T1) and after 10 (T10), 20 (T20), 28 (T28), and 53 (T53) days of exposure. All bars show the mean SSF ± standard errors from n = 2-3 bottle replicates of 2-3 pooled individuals. On days 10 and 20, n = 1 of 1-3 pooled individuals. The red lines or bars represent winter warming treatments (OW, 4 °C), and the blue lines or bars represent control temperature (0 °C). The dotted lines or bars with dashed marks indicate ocean acidification treatments (OA, pH = 7.4-7.3). Asterisks * indicates significant influence on the slope throughout the entire 53 days of the experiment (MLRM, \**P* < 0.05, \*\**P* < 0.01, \*\*\**P* < 0.001).

**Table 3.**
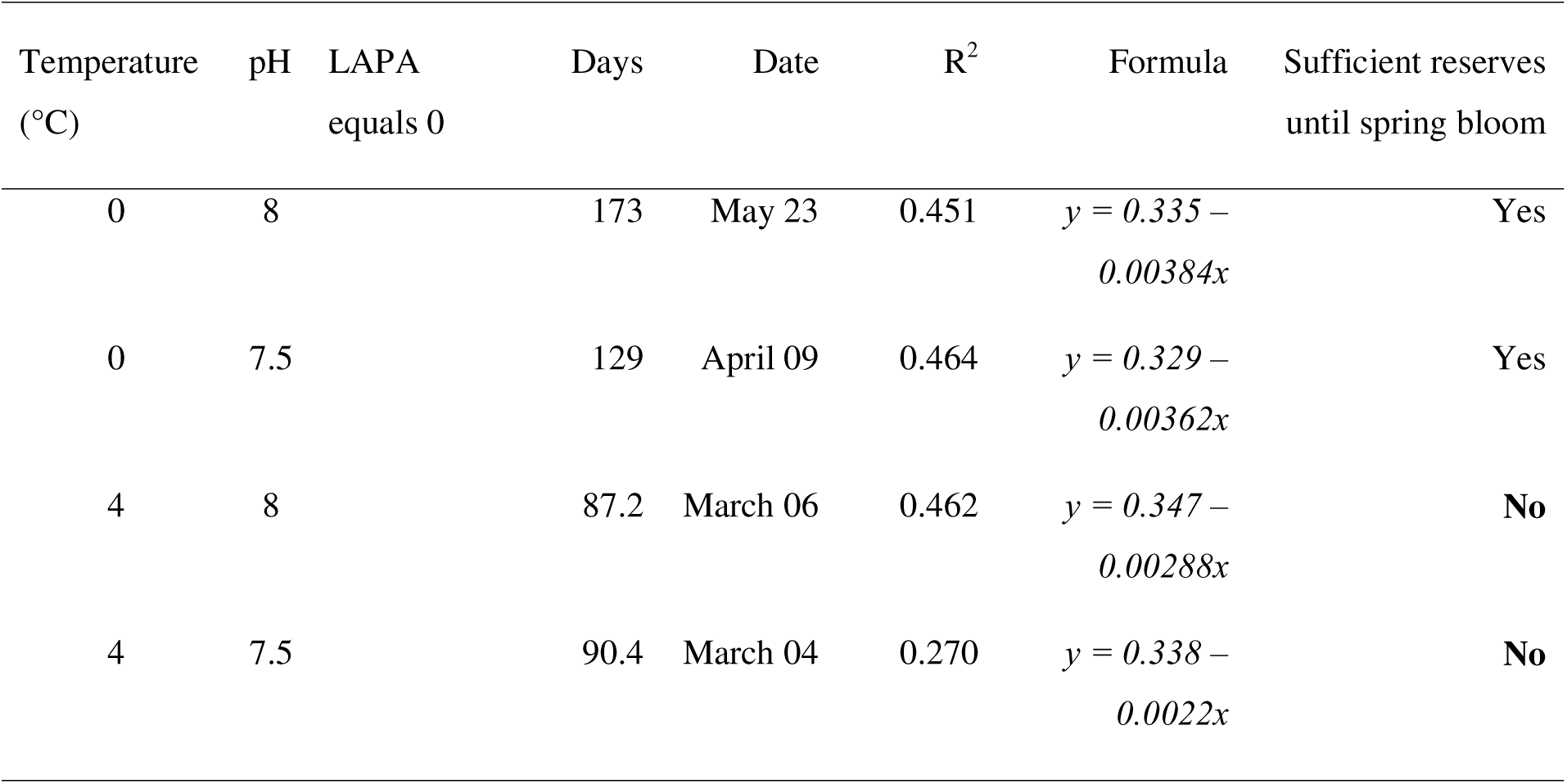
Expected time until 100% lipid depletion in *Calanus glacialis* within each treatment, predicted from linear regressions of each experimental treatment, not forcing the data through zero. Equations and R squared are presented in the table.

### 3.4 Oxygen consumption

OW significantly increased the oxygen consumption of *C. glacialis* (main effect of OW, Table 2, Figure 4c). Overall, OA had no significant main effect on the oxygen consumption of *C. glacialis*, but interactively modulated the OW effect. Specifically, OA reduced the oxygen consumption at 0°C, while increasing the oxygen consumption at 4°C (two-way ANOVA, Table 2, Figure 4c). This pH-temperature interaction was most pronounced (cumulative) between days 20 and 25 of the exposure period (Figure 4c), coinciding with the peak moulting period of copepodites into adults in the warming treatments (Figure 3b).

### 3.5 DNA damage

Both OA and OW individually and in combination caused DNA damage in *C. glacialis* (Figure 4d, Table S3 in Appendix S3). Acute exposure to OA, OW, or OWA resulted in an increasing DNA damage trend, with significant DNA damage observed in *C. glacialis* exposed to both stressors compared to controls (Fisher’s LSD at the 95% confidence level, *P* < 0.05, Table S3, Appendix S3). In the OA, the highest SSF values were observed after 53 days of continuous exposure.

### 3.6 Gene expression

Overall, there was no significant effect of temperature, OA, or the combined treatment on gene expression (see Table S1 for an overview of included genes, and Tables S4 and S5 for full statistical results); however, there were some patterns in the expression of within timepoints for some genes (Figure 5). Two genes, *ferritin* and *rad51,* were more expressed immediately after exposure compared with at the latter time point (28 days). Conversely, expression of *ercc1*, *parp1*, and *ogg1* was higher after 28 days in comparison to the acute response on day 1. The mean expression of *ferritin, parp1, ercc1, and ogg1* was higher under OW, and a similar but lower magnitude response to OA was observed. The expression of *ferritin, ercc1*, and *errcc4* showed a combined effect of OA and OW (Figure 5). Oxidative stress marker gene *ferritin* was significantly upregulated in the combined OW and OA treatment compared to warming alone after exposure for 28 days (Fisher’s LSD at the 95% confidence level, *P* < 0.05). The DNA repair genes *ercc1* and *ercc4* were significantly upregulated in the combined treatments compared to the control after 28 days of exposure (Fisher’s LSD at the 95% confidence level, *P* < 0.05).

**Figure 5.**
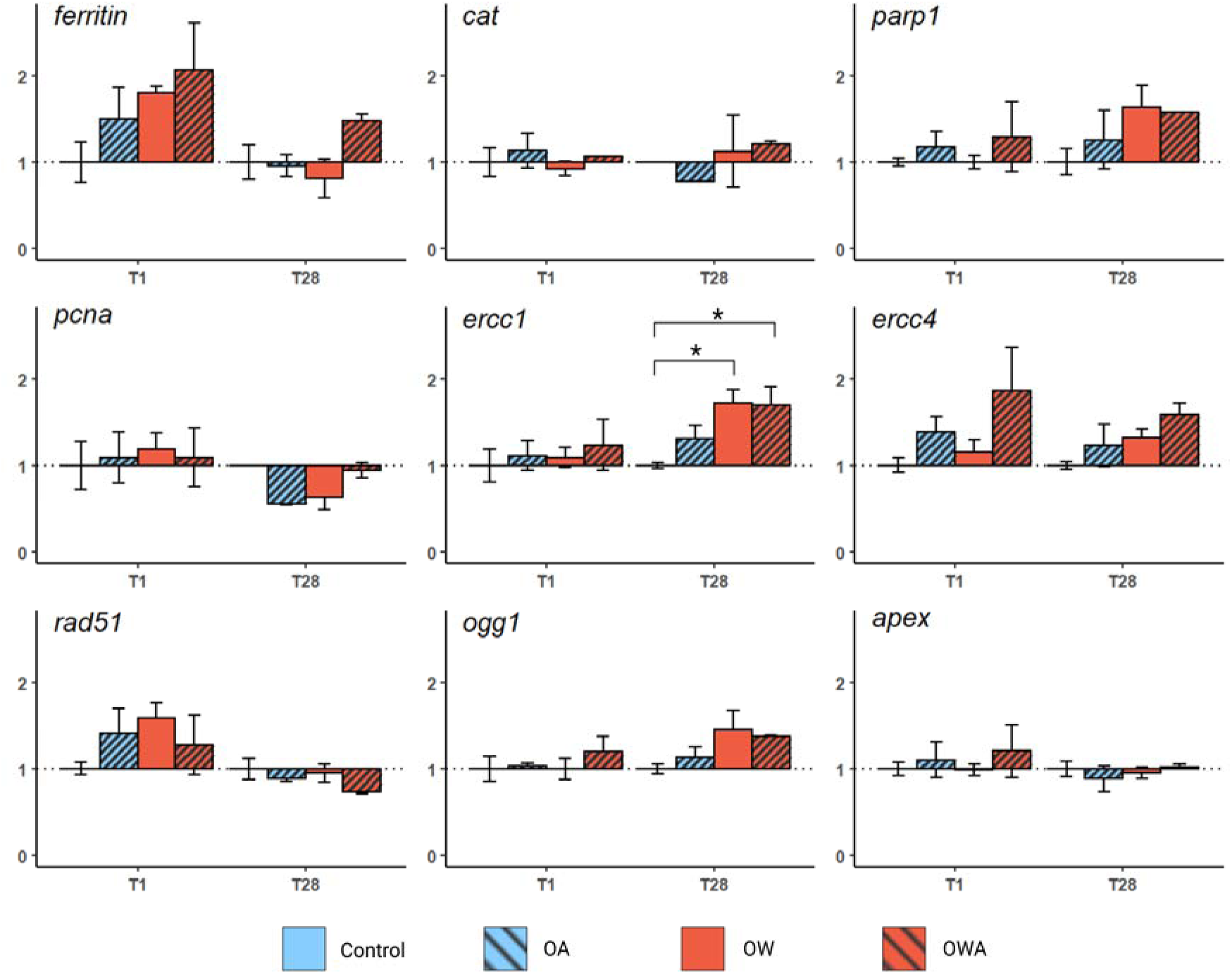
Relative gene expression (efficiency-adjusted ΔΔCP in *Calanus glacialis* for the selected genes *ferritin, cat, parp1, pcna, ercc1, ercc4, rad51, ogg1*, and, *apex* on days 1 (T1) and 28 (T28) of the experiment. The measured gene expression is shown here as the fold-change relative to the expression in the control treatment of that same gene at the same time point and relative to control gene expression. All bars show the mean gene expression ± standard deviation from n = 2-3 bottle replicates of 3-5 pooled individuals. Blue bars represent control temperature (0°C) treatments, and red bars show winter ocean warming (OW, 4°C) treatments. Bars with dashed marks represent ocean acidification (OA, pH = 7.4-7.3) treatments. Asterisks * indicates significant influence on the slope throughout the entire 53 days of the experiment (MLRM, \**P* < 0.05)

## 4. Discussion

Marine organisms are simultaneously exposed to multiple anthropogenic stressors, with climate change and ocean warming considered the most disruptive, causing cascading impacts throughout marine ecosystems (Alter et al., 2024; Dinh et al., 2023; Søreide et al., 2010). For the first time, our new findings on the combined effects of ocean warming and acidification on *Calanus glacialis*, a foundational Arctic marine grazer, uncover novel mechanisms by which OW and OA may synergistically disrupt overwintering during the Polar Night. These include increased winter mortality, accelerated development, heightened metabolism, and early lipid depletion, increased DNA damage, and low investment in maintenance and damage repair, creating a month-long phenological mismatch between overwintering survival, energy reserve, reproduction, and primary production. This mismatch underscores a severe bottleneck posed by the long, dark, and food-deprived winter, which can last up to six months in seasonally ice-covered waters in the high Arctic. These disruptions threaten the survival of marine grazers and may trigger cascading effects throughout Arctic marine ecosystems.

### 4.1 Warming disrupts the overwintering of Calanus glacialis

We found robust evidence that OW, as predicted by SOBH, disrupted the overwintering of *C. glacialis* by advancing the moulting from copepodite stage V to adults, and impairing metabolic depression, a state of significantly reduced metabolic rate, by increasing respiration, thereby depleting energy reserves. At 4°C, OW did not directly increase mortality but accelerated moulting over a month earlier than in the control at 0 °C, which falls within the winter temperature range -1.5 °C to +2.0 °C in Arctic shelves and fjords (Skogseth et al., 2020). This aligns with seasonal patterns observed in Arctic fjords, where moulting typically accelerates in late January (Hatlebakk et al., 2022). Early moulting, an energetically costly process, coincided with elevated oxygen consumption, particularly during peak moulting (December 20 - 28), reflecting disrupted metabolic depression critical for diapause (Baumgartner & Tarrant, 2017). Metabolic rates of diapausing copepods are generally about one-fifth of those for active copepods (Maps et al., 2014), and this process is governed by reduced anabolic enzyme activities (Freese et al., 2016). The heightened metabolic activities, also observed in other calanoid copepods under warming, such as *Acartia* sp. (Vehmaa et al., 2012), *Centropages tenuiremis* (Li & Gao, 2012), and the Arctic *C. hyperboreus* (Hildebrandt et al., 2014), is likely a result of increasing biosynthetic demands for moulting and gonad development (Karlsson & Søreide, 2024; Rey-Rassat et al., 2002).

Lipid reserves decreased faster in OW, which was associated with increased metabolism, likely fuelling moulting and gonad maturation (Freese et al., 2016). Our model showed that warming could result in a complete depletion of the lipid reserve by early March, which is at least one month before the first algal bloom in April in Billefjorden (Søreide et al., 2022) and Isfjorden (Vader et al., 2024) or elsewhere in the high Arctic (see Table S2 in Appendix S2, see also Leu et al., 2015). Even a new study showed that the photosynthesis of Arctic algae can happen as early as 14 March under extremely low light intensity (Hoppe et al., 2024), the first peak of polyunsaturated fatty acids (PUFAs) that are essential for the successful reproduction, growth, and development of *C. glacialis* females and nauplii only occurs in April-May due to low solar angle at extreme latitudes (Renaud et al., 2024; Søreide et al., 2010). A more than one-month asymmetry between the energy depletion of a foundational grazer like *C. glacialis* and the spring algal bloom is huge and critical, as the lipid reserve serves as the most important resource for early gonad maturation and reproduction of *C. glacialis* (Hatlebakk et al., 2022; Søreide et al., 2010).

The trend in elevated levels of DNA damage indicates possible acute genotoxic stress in the immediate onset of combined treatment (day 1), and chronic genotoxic stress at the end of exposure, which also seemed to follow earlier reported patterns in Arctic *A. longiremis* exposed to OA (Halsband et al., 2021) and is perhaps an indicator of indirect or oxidative stress-mediated genotoxicity, particularly after long-term exposure. Warming did not trigger major changes in the expression of genes relating to oxidative stress responses, including *ferritin* and *cat*. Immediate upregulation of *ferritin* does indicate some level of oxidative stress following acute exposure, which is in line with previous studies on this species (e.g., Smolina et al., 2015). This elevation persists in the combined treatment, indicating higher levels of stress. However, this pattern was not mirrored in the expression levels of *cat,* so it is not consistent with both oxidative stress marker genes. The observed higher metabolic activity within the OW treatments perhaps allowed for some DNA repair response, as indicated by the upregulation of *ercc1,* involved in DNA repair and corresponding low DNA damage after long-term thermal stress (Manandhar et al., 2015). This stands in contrast to *C. finmarchicus,* which is a North Atlantic species with increasing numbers in the European Arctic (Weydmann et al., 2018), a species that was shown to be much more responsive to thermal stress (Smolina et al., 2015) than *C. glacialis* in this study. A potential inability of *C. glacialis* to respond to their rapidly changing environment may make them highly vulnerable to a warming Arctic and to be replaced by boreal species, which are smaller, have less lipid, but are more adaptable to warming conditions like *C. finmarchicus* (Fossheim et al., 2015; Renaud et al., 2018; Weydmann et al., 2018). There is sufficient evidence of genotoxicity and some capacity for DNA repair in Arctic zooplankton exposed to climate-driven stressors (Bailey et al., 2017; Halsband et al., 2021), however, unpicking the dynamic mechanisms with different temporal profiles of response under different conditions, physiological states such as diapause, or altered metabolic levels during winter requires further investigation (Dinh et al., 2023; Halsband et al., 2021).

### 4.2 Beneficial effects of ocean acidification

Ocean acidification, with a pH of 7.4-7.3, alone had no observable negative effects on *C. glacialis*, yet it enhanced survival at the control temperature. Previous studies have often found a lack of OA impacts on Arctic marine species such as copepods (Bailey et al., 2017; Hildebrandt et al., 2016; Lewis et al., 2013, but see Thor et al., 2018) and fish (Dahlke et al., 2018; Dahlke et al., 2017). For example, at 0-1°C, Hildebrandt et al. (2014, 2016) found no changes in grazing, respiration rates, body mass, and mortality of *C. glacialis* females and copepodites exposed to a pH level of 7.2 over 16 to 62 days, from late summer (August) to autumn (November). In contrast, we find that OA had a beneficial effect on survival at low temperatures. Metabolic depression has been shown to be enhanced by OA in oysters, decreasing energy reserve consumption and increasing their survival (Lutier et al., 2022; 2025). Therefore, it is possible that, in our case, OA is beneficial to *C. galacialis* by interacting with metabolic depression that controls overwintering survival. This is the first observation of such a mechanism, which requires further research to be fully understood. DNA damage increased in OA and OW, and was highest in OWA and seemed to gradually increase over time in *C. glacialis* exposed to low pH at the control temperature, peaking after 53 days. It is possible that low pH directly affects the structure of DNA and reduces DNA repair (Lee et al., 2020; Lopes et al., 2019).

### 4.3 OW makes OA harmful for overwintering Calanus glacialis

The most striking and important finding in this study was the synergistic effects of OW and OA on the survival of *C. glacialis*. Indeed, while at 0°C OA was beneficial for survival (see part 4.2), it drastically increased mortality at 4°C for *C. glacialis*, as predicted by SOBH. This mirrors the OWA synergistic effect observed in Arctic fish such as Atlantic and polar cods (Dahlke et al., 2018), pteropods (Lischka & Riebesell, 2012), and temperate copepod *Acartia tonsa* (Dam et al., 2021). In our case, the synergistic effect could be induced by OW, causing the organism to exit metabolic depression (see part 4.1), thereby removing the protective effects of this phenomenon under OA. Typically, *C. glacialis* overwinters in deep Arctic waters at temperatures ranging from -1.8 °C to 2 °C, where metabolic depression, characterised by reduced respiration and anabolic activity, is critical for energy conservation and survival (Baumgartner & Tarrant, 2017; Maps et al., 2014). At these low temperatures, *C. glacialis* maintains a low internal pH without physiological impairment (pH as low as 5.5, Freese et al., 2015). Indeed, metabolic depression minimises the cost of acid-based regulation, allowing for saving energy reserves and extending survival (Pörtner & Bock, 2000; Pörtner et al., 2000; Storey & Storey, 1990). However, at 4 °C, *Calanus* exits from metabolic depression, and this metabolic shift likely abolishes the protective effects of diapause against OA. Here, we show that diapause during the Polar Night regulates the survival of overwintering copepods exposed to OA. By affecting diapause, OW could drastically increase mortalities during this crucial time for moulting and reproduction, representing a bottleneck in the life cycle and thus population recruitment.

### 4.4 Stressed Overwintering Bottleneck Hypothesis (SOBH) and applications beyond the Arctic marine ecosystems

The overwintering of Arctic zooplankton is linked to one of the largest animal vertical migrations on Earth, which can profoundly affect the food web structure (Bandara et al., 2021), the lipid and pollutant transfer between trophic levels (Borgå et al., 2022; Årthun et al., 2025), and the carbon cycle of the Arctic seas and ocean (Jonasdottir et al., 2015). However, deep waters in the Arctic, such as fjords in Svalbard, the Barents Sea, and Disko Bay in West Greenland, where *C. glacialis* populations overwinter, have been increasingly impacted by warm Atlantic water (Polyakov et al., 2017; Årthun et al., 2025). Indeed, temperatures in deep waters have risen by 1 - 4 °C over the past two to three decades (Lischka & Riebesell, 2012; Myers & Ribergaard, 2013; Skogseth et al., 2020), signalling that overwintering disruption is already underway during the warming Polar Night, making copepods more vulnerable to OA. Such overwintering disruption potentially contributes to observed rapid shifts in *Calanus* distributions (Feng et al., 2018; Kvile et al., 2018) by reducing winter survival, impairing post-winter reproduction, and the Arctic biological pump efficiency (up to -40% by 2100, Oziel et al., 2025). Notably, these shifts have not been fully explained by thermal adaptation (Hinder et al., 2014), sea ice changes, or timing and duration of spring blooms (Feng et al., 2018). Our results, as predicted by SOBH, uncover hidden mechanisms by which OA amplifies OW-induced disruption in key overwintering Arctic zooplankton grazers like *C. glacialis*, thereby contributing to broader ecological shifts and emphasizing overwintering disruption as an underappreciated driver of change. These impacts can be more severe in the future, especially for calcifying organisms, e.g., Arctic pteropods (Lischka & Riebesell, 2012).

At a broader scale, Arctic ecosystems may be approaching critical tipping points (Heinze et al., 2021), with eight out of sixteen global climate tipping points located within the Arctic Circle (Armstrong McKay et al., 2022; Lenton et al., 2023). SOBH-famed findings of AO-amplified overwintering disruption in *Calanus* copepods could hasten these transitions by widening the gap of phenological mismatches between the secondary producer with primary producers and marine predators (Figure 6), thereby reducing top-down control of algal blooms and the bottom-up effects on the transfer of energy, nutrients (e.g., PUFAs), and pollutants to higher trophic levels, such as Arctic fish (Grosbois et al., 2022), birds (e.g., *Alle alle*, Karnovsky et al., 2003), and mammals (Årthun et al., 2025).

**Figure 6.**
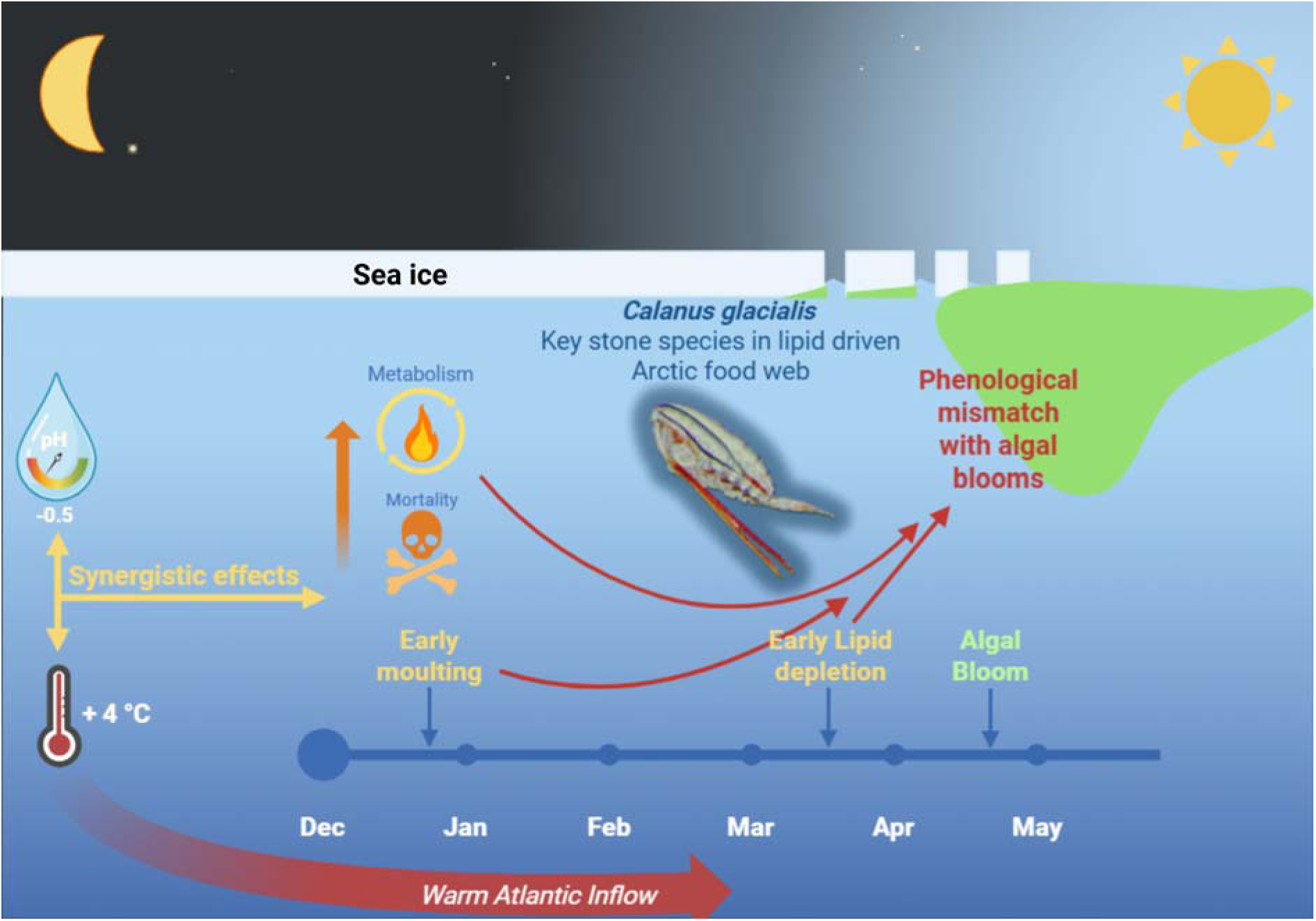
Stressed Overwintering Bottleneck Hypothesis (SOBH): Conceptual pathways in which winter ocean warming and ocean acidification can result in overwintering disruption through drastically increased winter mortality, accelerated moulting, heightened metabolism, and early lipid depletion, ultimately generating month-long phenological mismatches among overwintering survival, energy reserves, reproduction, and primary production.

The SOBH can be applied beyond OWA interaction, positioning overwintering as a universal stressor-vulnerability window across ecosystems, reshaping Arctic and global winter biodiversity (Dinh et al., 2023). For example, Arthropods are the most diverse group with over 80% of all identified animal species (∼1.17 million species, Ødegaard, 2000) and 50% of total animal biomass (1 Gt carbon equivalents, Bar-On et al., 2018), and the majority of these animals can overwinter in different forms of dormancy (Baumgartner & Tarrant, 2017; Hopkin, 1997; Koštál et al., 2017; Potapov et al., 2023), which are increasingly impacted by various types of anthropogenic stressors (Dinh et al., 2023). In marine ecosystems, pollutants like pyrene can amplify toxicity 300-fold in overwintering *C. glacialis* (Toxværd et al., 2018), which can be further impacted by warming and microplastics (Albini et al., 2024).

In freshwater, shallow, winter-ice-covered lakes, hypoxia can be detrimental to invertebrates and fish (Jansen et al., 2025). While winter warming can delay the timing of ice cover in lakes, thereby promoting winter-active consumers by extending access to high-quality fat reserves from algae in early winter, this may strongly affect winter-inactive and benthic invertebrates in the spring, reshaping food web dynamics (Hébert et al., 2021). However, this study assumes that cyclopoid copepods can maintain low metabolism during the winter (Hébert et al., 2021). However, we show that the warming-induced increase in metabolism in *Calanus* copepods occurs during the middle of winter, when food is not available, which is a critical factor to determine the depletion of energy reserves, survival, and post-winter reproduction. Elevated metabolic rate is a universal response of organisms to warming, and Arctic species show the greatest warming-induced relative metabolic changes (Dillon et al., 2010). In terrestrial ecosystems, UK long-term (40 years) data revealed that warmer and wetter winters widen the seasonal mismatch between insectivorous birds (e.g., blue tits, *Cyanistes caeruleus*) and their food source, the winter moth (*Operophtera brumata*), risking extirpation if breeding lags > 24 days past peak prey (Simmonds et al., 2020). Future research must prioritise overwintering multiple-stressor experiments across multiple trophic levels and the ecosystem changes to refine predictive models, guide the Intergovernmental Panel on Climate Change (IPCC) forecasts for polar resilience amid tipping points (Lee et al., 2023) and inform winter conservation strategies for Arctic and global ecosystems facing unprecedented changes.

## Author contributions

J.E.S., K.V.D., J.D., M.L., and H.R. conceived the first idea for the study and discussed it with all co-authors. All co-authors contributed to the idea. J.D., M.L., L.S, J.S. and N.T. conducted the experiment, collected and analysed the data. J.D. and K.V.D. co-wrote the first version of the manuscript. All co-authors edited and approved the final manuscript.

## Acknowledgements

We would like to thank Michael Lemke, Anna Miettinen and Marta Paludetto for assistance during parts of the experiment, Henningsen Transport and guiding for their sampling support with MS Farm, and Sabine Marty, NIVA, for providing the carbonate chemistry analyses. This study was financially funded by FRAM – High North Research Centre for Climate and the Environment (project CLEAN - Cumulative impact and risk associated with multiple stressors in High North), Young Research Talent grant (RCN #325334), and the Nansen Legacy (RCN #276730).

## Data availability statement

The data that support the findings of this study are openly available in Figshare repository at DOI 10.6084/m9.figshare.30285304

## Ethics statement

We followed the standard operating procedure for conducting experiments approved by The University Centre in Svalbard, the University of Oslo, and the Research Council of Norway. Copepod *Calanus glacialis* used in this study is not classified as a protected invertebrate under the Norwegian Regulation on Animal experimentation (Forskrift om bruk av dyr i forsøk) and the European guidelines (Directive 2010/63/EU). Therefore, this research does not require formal ethical approval from an institutional committee on animal care and use. Zooplankton care, maintenance, and husbandry followed routine institutional protocols. All authors have seen and agreed to submit this version of the manuscript to *Global Change Biology Communications*.

