## Appendix S1 for "Stressed Overwintering Bottleneck Hypothesis: Ocean warming and acidification synergistically disrupt Arctic zooplankton overwintering"

Running title: Multistress-induced overwintering disruption

Jildou Dijkstra[1,2], Luise Schott[3], Nele Thomsen[4], Helena Reinardy[2,4], Mathieu Lutier[5], Janne E. Søreide[2,*], Khuong V. Dinh[5,*]

[1] Freshwater and Marine Ecology (FAME), University of Amsterdam, P.O. Box 94248, 1090 GE Amsterdam, The Netherlands

[2] The University Centre in Svalbard (UNIS), P.O. Box 156, Longyearbyen, Norway

[3] Freie Universität Berlin, Institut für Biologie, Altensteinstr. 6, D-14195, Berlin, Germany

[4] Scottish Association for Marine Science, PA37 1QA, Oban, Scotland

[5] Section for Aquatic Biology and Toxicology, Department of Biosciences, University of Oslo, Blindernveien 31, 0371 Oslo, Norway

### Appendix S1

### Supplementary methods

*Temperature and light logger data*

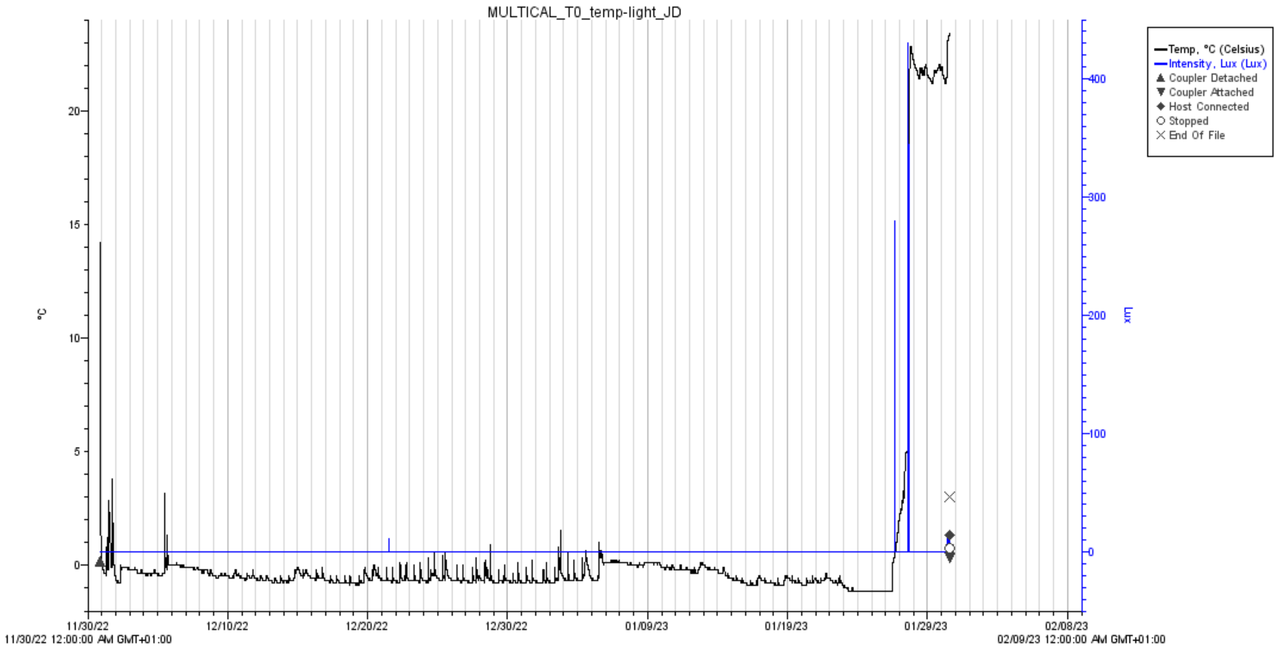


**Figure S1**. Data of temperature and light intensity measured every 5 minutes using a HOBO® Pendant (Onset, Bourne, USA) temperature and light logger (UA-002-08) under controlled conditions at 0 °C.

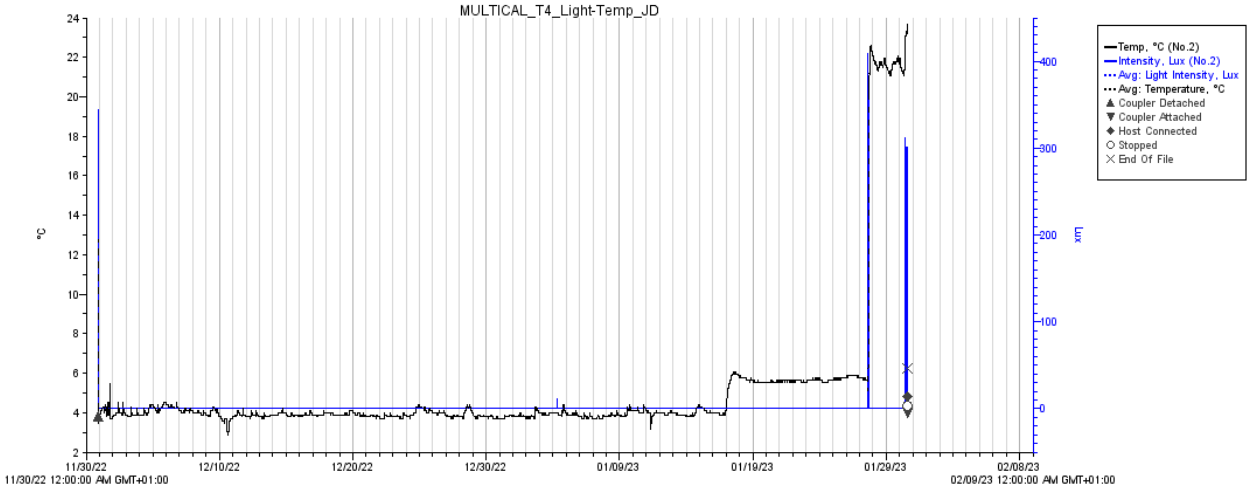


**Figure S2**. Data of temperature and light intensity measured every 5 minutes using a HOBO® Pendant (Onset, Bourne, USA) temperature and light logger (UA-002-08) under controlled conditions at 4 °C.

*Oxygen consumption*

Twice daily dissolved oxygen levels were measured at 09:00 and 18:00, ensuring levels never dropped below 60% saturation throughout the entire incubation. Oxygen levels were determined using a fibre optic cable connected to the OXY-4 4-Channel Fiber-Optic Oxygen Meter (PreSensTM, GmbH, Regensburg, Germany). On the inner surface of these bottles, a SP-PSt3 mini oxygen sensor (Loligo® Systems, Edinburgh, Scotland; accuracy ± 0.4% $O_2$ at 20.9% $O_2$) was attached. Sensor spots were calibrated prior to and directly after the experiment, using 100% oxygenated seawater and a 0% oxygen solution of sodium sulphite in demineralized water (1gr: 100 mL $Na_2SO_3$:MilliQ), to account for potential drift in sensor sensitivity throughout the experiment. Oxygen levels were measured in 12 bottles per treatment and in all controls between days 1 and 28. After day 28, oxygen levels were measured in 6 bottles per treatment. For the time interval between water changes, the oxygen consumption per bottle was calculated as the slope of the regression line of the oxygen concentration and time (Morata & Søreide, 2015). From this, the individual oxygen consumption per individual, corrected for dry weight, was determined by dividing total consumption by the number of alive individuals present in each bottle. These values were corrected for any other seawater background metabolic activities using the oxygen consumption measured in three blanks (only seawater) for each corresponding treatment. Respiration rates were then converted to carbon demand using a respiration conversion factor of 1 mol $O_2$: 0.97 mol $CO_2$ (Hernández-León & Ikeda, 2005).

*RNA extraction and cDNA synthesis for quantitative reverse transcriptase PCR*

Surviving zooplankton for DNA damage and gene expression analyses were fixed in 2 mL cryotubes in RNA*later* stabilization solution (Thermo Fisher Scientific, Waltham, MA, USA) (5 : 1 RNA*later* : FSW) immediately after being removed from the experimental bottle. Individuals were stored at -80°C for one month before transportation on dry ice to the Scottish Association for Marine Science (SAMS) for further analysis.

RNA was extracted with the Qiagen RNeasy Mini Kit following the manufacturers protocol with adjustments for chitinous copepods. A pool of 3 to 5 copepods was homogenized stepwise with handheld micropestles in 50, 150, and 350 µl of RLT buffer to effectively break open the exoskeleton. The homogenate was centrifuged for 1 min at 8,000 *g* to remove large remnants of the exoskeleton, and the supernatant was transferred to a new microcentrifuge tube. After adding 350 µl of 70% ethanol, the solution was transferred to 2 ml spin columns. In between two washing steps, a 15-minute DNA digestion in 80 µl DNase (10 µl DNase

I, 70 µl RDD buffer) was performed at room temperature. Two final washing steps in 500 µl RLT buffer were followed by centrifugation for 2 min at 8,000 *g* and 1 min at 14,000 *g* to fully expel remnants of the buffer from the column membrane. Lukewarm nuclease-free water was added to the spin column membrane and left at room temperature for 20 minutes, after which total RNA was eluted by 1 minute centrifugation at 8,000 *g*. RNA quantity and quality were determined on a Nanodrop (ThermoFisher Scientific Inc., Massachusetts, US) and verified on a 0.7% agarose gel (1x TAE buffer, 1 h at 75 V). Low quality RNA (260:230 ratio < 1.7) was cleaned up with the Qiagen RNeasy PowerClean Pro Clean-up Kit. 500 ng total RNA was reverse transcribed with the High-Capacity cDNA Reverse Transcription Kit (Applied Biosystems) and stored at -20 °C.

*Quantitative reverse transcriptase PCR*

Nine genes were selected for their known function in either the oxidative stress response (*ferritin* (Roncalli et al., 2023), *cat* (Hansen et al., 2008) or DNA repair system (*ercc1* (Chipchase & Melton, 2002; Rhee et al., 2012), *ercc4* (Gan et al., 2020; Gregg et al., 2011), *pcna* (Reinardy & Bodnar, 2015; Rhee et al., 2012), *parp1* (Buendia-Padilla et al., 2023; Reinardy & Bodnar, 2015; Sahlmann et al., 2019), *apex* (Jayaraman et al., 1997; Meira et al., 2001), *ogg1* (Sahlmann et al., 2019), *rad51* (Fasullo et al., 2001; Rapp & Greulich, 2004; Rhee et al., 2012). Primers were extracted from literature or designed from published transcriptome sequences (Table S1). Primers were designed for a product length of 100 – 150 bp and checked for hairpin structures, complementarity, secondary structures, and self-annealing with Primer3Plus (Untergasser et al., 2012) and BLAST from the National Center for Biotechnology Information (NCBI). Primer pairs were verified for qPCR product length and primer specificity on a 1.5% agarose gel (1x TAE buffer, 1 h 15 min at 100 V). Differential expression of selected genes was analysed by qRT-PCR with a SYBR Green detection method (Reinardy and Bodnar, 2013) on the Lightcycler96 (Rocher Diagnostics Corporation, USA). The PCR protocol consisted of preincubation (95 °C for 600s), three-step amplification (95 °C for 15s, 56 °C for 60s, 72 °C for 60s), and melt curve analysis (95 °C for 15s, 60 °C for 60s, 97 °C for 1s). A 5-fold standard curve for each primer pair was run on the same plate as the quantified samples (triplicates), and qPCR efficiency was checked to be between 90 and 110%. Fold change was calculated according to the efficiency-adjusted ΔΔCP method by Pfaffl (2001), relative to treatment controls and the geometric mean of three control genes (*β-actin*, *gapdh*, and *efa1*α).

**Table S1**. Genes targeted and corresponding primers used for q-RT-PCR.

| Gene function | Gene | Gene pathway | GenBank Accession No. | Primer | Primer Sequence (5'-3') | Product length (bp) | Reference |
|---|---|---|---|---|---|---|---|
| Control | *β-actin* | | ES414833 | F<br>R | CAACCTTCTTGCAGCTCCTCCG<br>CCCACGATGGAGGGGAAGACGG | 124 | Hansen et al., 2007 |
| | *ef1a* | | GBXU01018644 | F<br>R | AGTGGAGGCCAGCTCAAACAT<br>ACGGGTCTGACAGTGGTTTCC | 143 | Present study |
| | *gapdh* | | HQ270535 | F<br>R | ATCTTGAAGGGTGGTGCCAAG<br>GTCTTCTGAGTGGCAGTGATGG | 126 | Lauritano et al., 2012 |
| Oxidative stress | *ferritin* | Metal ion homeostasis | GH271794 | F<br>R | GTCACTGGGCATGTCAATGG<br>CAGAGAGGAGAAAGGAAGCGG | 129 | Hansen et al., 2010 |
| | *cat* | ROS detoxification | EL965956 | F<br>R | CATGGTCAGCAGGCAAAGAA<br>CATGGTCAGCAGGCAAAGAA | 142 | Present study |
| DNA repair | *ercc1* | Nucleotide excision repair | GBXU01006032 | F<br>R | TGGACAGCAGGACATGGACTT<br>ACTGGAGATGGCGAGAAAGGC | 146 | Present study |
| | *ercc4* | Nucleotide excision repair and DNA recombination | GBXU01005979 | F<br>R | TGTGCAGCCTGAATGTTGG<br>TGTGCAGCCTGAATGTTGG | 116 | Present study |
| | *pcna* | Base excision repair | GBXU01009868 | F<br>R | GGACAGCGTGGGCAAGGTT<br>GGCAGCACCGATCACCACTA | 125 | Present study |
| | *parp1* | Base excision repair | GBXU01009226 | F<br>R | GCAGCCTTGGCTTCTGGAAC<br>TGTCATCCAGCGTCGTCGAC | 170 | Present study |
| | *apex* | Base excision repair | GBXU01024194 | F<br>R | TGAAGGAGCGGATCGTGGAT<br>GGGAAGGGTCTCCACACTCC | 148 | Present study |
| | *ogg1* | Base excision repair | GBXU01024371 | F<br>R | ATCGGTGCAGGAAACTGAGC<br>CACAGCAGCCACCAAGGAAT | 116 | Present study |
| | *rad51* | Homologous recombination | GBXU01013784 | F<br>R | AATGGCTGGTGAGGAAGGTG<br>ATCACGCCGATAGACAGGGT | 138 | Present study |

*DNA damage*

The fast micromethod assay (FMM) for the detection of DNA strand breaks was conducted on individual copepods to quantify DNA damage. The method described by Halsband et al. (2021) was followed with adjustments for detection in individual copepods. Replicate copepods were taken from sample bottles and individually homogenized in 50 µl homogenization buffer (20 mM EDTA, 10% DMSO), and total volume was adjusted to 260 µl to dilute the chitinous exoskeleton. 20 µl homogenate was pipetted into quadruplicate wells of a black flat bottom 96-well plate and lysed for 40 min in the dark (20 µl 9 M urea, 0.1% SDS, 0.2 M EDTA, 2% Quant-iT ™ PicoGreen®). An unwinding solution (200 µl, 20mM EDTA, 1M NaOH, pH adjusted to pH 13.5) was added after lysis and fluorescence readings taken immediately (POLARstar Omega, BMG Labtech Ltd; gain: 1500, excitation: 485, emission: 520) followed by every 5 min for 30 min. Strand scission factors (SSFs) that quantify relative DNA strand breaks were calculated after 20 min according to Schröder et al. (2006). Treatment SSFs are relative to controls within their respective time point.
