## Appendix S2 for "Stressed Overwintering Bottleneck Hypothesis: Ocean warming and acidification synergistically disrupt Arctic zooplankton overwintering"

Running title: Multistress-induced overwintering disruption

Jildou Dijkstra[1,2], Luise Schott[3], Nele Thomsen[4], Helena Reinardy[2,4], Mathieu Lutier[5], Janne E. Søreide[2,*], Khuong V. Dinh[5,*]

[1] Freshwater and Marine Ecology (FAME), University of Amsterdam, P.O. Box 94248, 1090 GE Amsterdam, The Netherlands

[2] The University Centre in Svalbard (UNIS), P.O. Box 156, Longyearbyen, Norway

[3] Freie Universität Berlin, Institut für Biologie, Altensteinstr. 6, D-14195, Berlin, Germany

[4] Scottish Association for Marine Science, PA37 1QA, Oban, Scotland

[5] Section for Aquatic Biology and Toxicology, Department of Biosciences, University of Oslo, Blindernveien 31, 0371 Oslo, Norway

### Appendix S2. Timing of the Arctic algal blooms

Table S2: Timing of Arctic Algal Blooms

| Location | Coordinates (Lat, Lon) | Timing – Now | References | Timing - Future projection | References |
|---|---|---|---|---|---|
| Isfjorden (Spitsbergen, Svalbard) | 78.2° N, 15.0° E | Late April - June | Hoppe et al., 2024; Vader et al., 2024 | Earlier bloom due to Atlantification ~5 days earlier per decade | Henson et al., 2018; Knies et al., 2025; Payne & Roesler, 2019 |
| Kongsfjorden (Ny-Ålesund region, Svalbard) | 78.9° N, 11.9° E | Late April - May | Assmy et al., 2023; Hegseth & Tverberg, 2013 | Earlier bloom due to Atlantification ~5 days earlier per decade | Henson et al., 2018; Knies et al., 2025; Payne & Roesler, 2019 |
| Barents Sea | 74–80° N, 15–40° E | April - May | Nicholson et al., 2025; Oziel et al., 2017 | 1 month earlier | Oziel et al., 2017; Årthun et al., 2025 |
| Disko Bay (Qeqertarsuup Tunua, W Greenland) | 69° N, 52° W | April - June | Dunweber et al., 2010 | Earlier bloom due to low ice cover and enhanced autumn bloom due to glacial melt | Møller et al., 2023; Wood et al., 2025 |
| Young Sound (Young Sund, NE Greenland) | 74.3° N, 20.3° W | July - October | Maar et al., 2025 | Longer bloom window | Maar et al., 2025 |
