## Appendix S3 for "Stressed Overwintering Bottleneck Hypothesis: Ocean warming and acidification synergistically disrupt Arctic zooplankton overwintering"

Running title: Multistress-induced overwintering disruption

Jildou Dijkstra[1,2], Luise Schott[3], Nele Thomsen[4], Helena Reinardy[2,4], Mathieu Lutier[5], Janne E. Søreide[2,*], Khuong V. Dinh[5,*]

[1] Freshwater and Marine Ecology (FAME), University of Amsterdam, P.O. Box 94248, 1090 GE Amsterdam, The Netherlands

[2] The University Centre in Svalbard (UNIS), P.O. Box 156, Longyearbyen, Norway

[3] Freie Universität Berlin, Institut für Biologie, Altensteinstr. 6, D-14195, Berlin, Germany

[4] Scottish Association for Marine Science, PA37 1QA, Oban, Scotland

[5] Section for Aquatic Biology and Toxicology, Department of Biosciences, University of Oslo, Blindernveien 31, 0371 Oslo, Norway

**Appendix 3**

**Table S3**: Statistical Summary of DNA damage (SSF) one-way ANOVA

| Timepoint | Test | Factor | p-value | Post hoc |
|---|---|---|---|---|
| T1 | One-way ANOVA | treatment | 0.106 | *** Fisher LSD**<br>**Control & OWA** |
| T28 | One-way ANOVA | treatment | 0.0739 | *** Fisher LSD**<br>**OA & OWA** |
| T53 | One-way ANOVA | treatment | 0.007 | *** Fisher LSD**<br>**OA & OW**<br>**OA & OWA** |

**Table S4**. Statistical Summary of Gene Expression One-way ANOVA.

| Gene | Timepoint | Test used | Factor | F statistic (df) | p-value | Post-hoc notes |
|---|---|---|---|---|---|---|
| *ferritin* | T1 | One-way ANOVA | treatment | F(3,19) = 2.12 | 0.242 | |
| | T28 | One-way ANOVA | treatment | F(3,19) = 2.12 | 0.191 | |
| *cat* | T1 | One-way ANOVA | treatment | F(3,12) = 0.20 | 0.897 | |
| | T28 | One-way ANOVA | treatment | F(3,12) = 0.20 | 0.936 | |
| *parp1* | T1 | One-way ANOVA | treatment | F(3,18) = 0.78 | 0.933 | |
| | T28 | One-way ANOVA | treatment | F(3,18) = 0.78 | 0.421 | |

| | | | | | | |
|---|---|---|---|---|---|---|
| *pcna* | T1 | One-way ANOVA | treatment | F(3,16) = 0.15 | 0.971 | |
| | T28 | One-way ANOVA | treatment | F(3,16) = 0.15 | 0.217 | |
| *ercc1* | T1 | One-way ANOVA | treatment | F(3,18) = 2.42 | 0.902 | |
| | T28 | One-way ANOVA | treatment | F(3,18) = 2.42 | **0.065** | *** Fisher LSD OW > Control OWA > Control** |
| *ercc4* | T1 | One-way ANOVA | treatment | F(3,19) = 4.11 | 0.113 | |
| | T28 | One-way ANOVA | treatment | F(3,19) = 4.11 | 0.200 | |
| *rad51* | T1 | One-way ANOVA | treatment | F(3,19) = 0.54 | 0.423 | – |
| | T28 | One-way ANOVA | treatment | F(3,19) = 0.54 | 0.383 | – |
| *ogg1* | T1 | One-way ANOVA | treatment | F(3,19) = 1.42 | 0.688 | – |
| | T28 | Kruskal-Wallis | treatment | F(3,19) = 1.42 | 0.127 | – |
| *apex* | T1 | One-way ANOVA | treatment | F(3,18) = 0.40 | 0.862 | – |
| | T28 | One-way ANOVA | treatment | F(3,18) = 0.40 | 0.783 | – |

**Table S5**. Statistical Summary of Gene Expression two-way ANOVA.

| Gene | Treatment p-value | Timepoint p-value | Interaction p-value | Overall significant? | Significant post-hoc results |
|---|---|---|---|---|---|

| | | | | | |
|---|---|---|---|---|---|
| *ferritin* | 0.088 | 0.19 | 0.407 | No | |
| *cat* | 0.953 | 0.652 | 0.837 | No | |
| *parp1* | 0.793 | 0.74 | 0.59 | No | |
| *pcna* | 0.954 | 0.359 | 0.693 | No | |
| *ercc1* | 0.856 | 0.979 | 0.371 | No | |
| *ercc4* | **0.075** | 0.655 | 0.801 | No | ***Tukey HSD: treatment T4P75 > T0P8 (p=0.028)*** |
| *rad51* | **0.204** | 0.081 | 0.362 | No | ***Tukey HSD: timepoint T28<T1 (p=0.011)*** |
| *ogg1* | 0.706 | 0.635 | 0.421 | No | |
| *apex* | 0.756 | 0.627 | 0.887 | No | |
